# A detailed behavioral, videographic, and neural dataset on object recognition in mice

**DOI:** 10.1101/2022.05.10.491259

**Authors:** Chris C. Rodgers

## Abstract

Mice adeptly use their whiskers to touch, recognize, and learn about objects in their environment. This ability is enabled by computations performed by populations of neurons in the somatosensory cortex. To understand these computations, we trained mice to use their whiskers to recognize different shapes while we recorded activity in the barrel cortex, which processes whisker input. Here, we present a large dataset of high-speed video of the whiskers, along with rigorous tracking of the entire extent of multiple whiskers and every contact they made on the shape. We used spike sorting to identify individual neurons, which responded with precise timing to whisker contacts and motion. These data will be useful for understanding the behavioral strategies mice use to identify objects by touch, as well as the neuronal responses that mediate those strategies. More generally, our carefully curated labeled data could be used to develop new computer vision algorithms for tracking body posture, or for extracting responses of individual neurons from large-scale neural recordings.

## Background & Summary

Active sensing is the process of moving the body in order to learn about the world^1^. For instance, animals can direct their eyes and ears toward an object of interest^2^. Rodents like mice have evolved sophisticated capabilities to move their whiskers onto objects in order to identify them, which enables them to navigate their world and interact with conspecifics^3^. This active touch is similar to the way primates are able to recognize objects by touching them with their hands and fingers^4,5^. While sensory processing and motor control are often studied in isolation, a fundamental challenge in systems neuroscience is to understand how perception and action work together to support natural behavior.

To study this, we previously^6^ trained head-fixed mice to discriminate concave from convex shapes using only their whiskers (hereafter, “shape discrimination”). As a control, another group of mice learned simply to detect the presence of either shape, versus catch trials when no shape was presented (“shape detection”). As mice did this, we used high-speed videography to precisely monitor the position of the whiskers and every contact that they made on the shape. This behavioral data provides a detailed picture of the motor exploration strategies mice used to perform each task, and the sensory input that they collected and used to make their decision.

Neural circuitry in the brain controls this motor exploration and processes the resulting sensory input. We recorded neural activity from the barrel cortex^7^, which processes somatosensory input from the whiskers, as mice performed the shape discrimination and detection tasks. Using a high-density extracellular recording array, we recorded neurons simultaneously across all layers of the barrel cortex, as well as the local field potential (LFP).

We previously used these data to identify how mice discriminated and detected shapes^6^. We found that shape discrimination mice compared the prevalence of contacts across whiskers, a comparison computation enabled by tuning changes in the barrel cortex. In contrast, shape detection mice simply responded to the total number of contacts across all whiskers, and in that case barrel cortex neurons maintained their expected tuning. We have also reported that barrel cortex represents sensory and motor variables in approximately orthogonal subspaces, which could be useful for generalization to novel conditions^8^. This flexible neural representation of the body and world could be useful for the active touch behaviors that mice evolved to do, which are more complex than the ones we study in the lab.

Here, we present the entire behavioral, video, and neural datasets that we collected during these tasks, available at https://dandiarchive.org/dandiset/000231. The dataset includes 88.9 hours of behavioral video at 200 frames per second, and 33.8 hours of 64-channel neural recording at 30 kHz. We provide these data both “raw”—as we acquired them—and also in a processed, aligned, and documented format using the Neurodata Without Borders (NWB) standard (Figure 1)^9^. This processing primarily consisted of two steps: we used a deep neural network to track the whiskers in video and we used spike sorting to identify individual neurons in the recordings. This dataset improves on our original data release^10^ because it includes more raw data, including videos and full bandwidth recordings, and it is now formatted in the more convenient and complete NWB standard.

**Figure 1.**
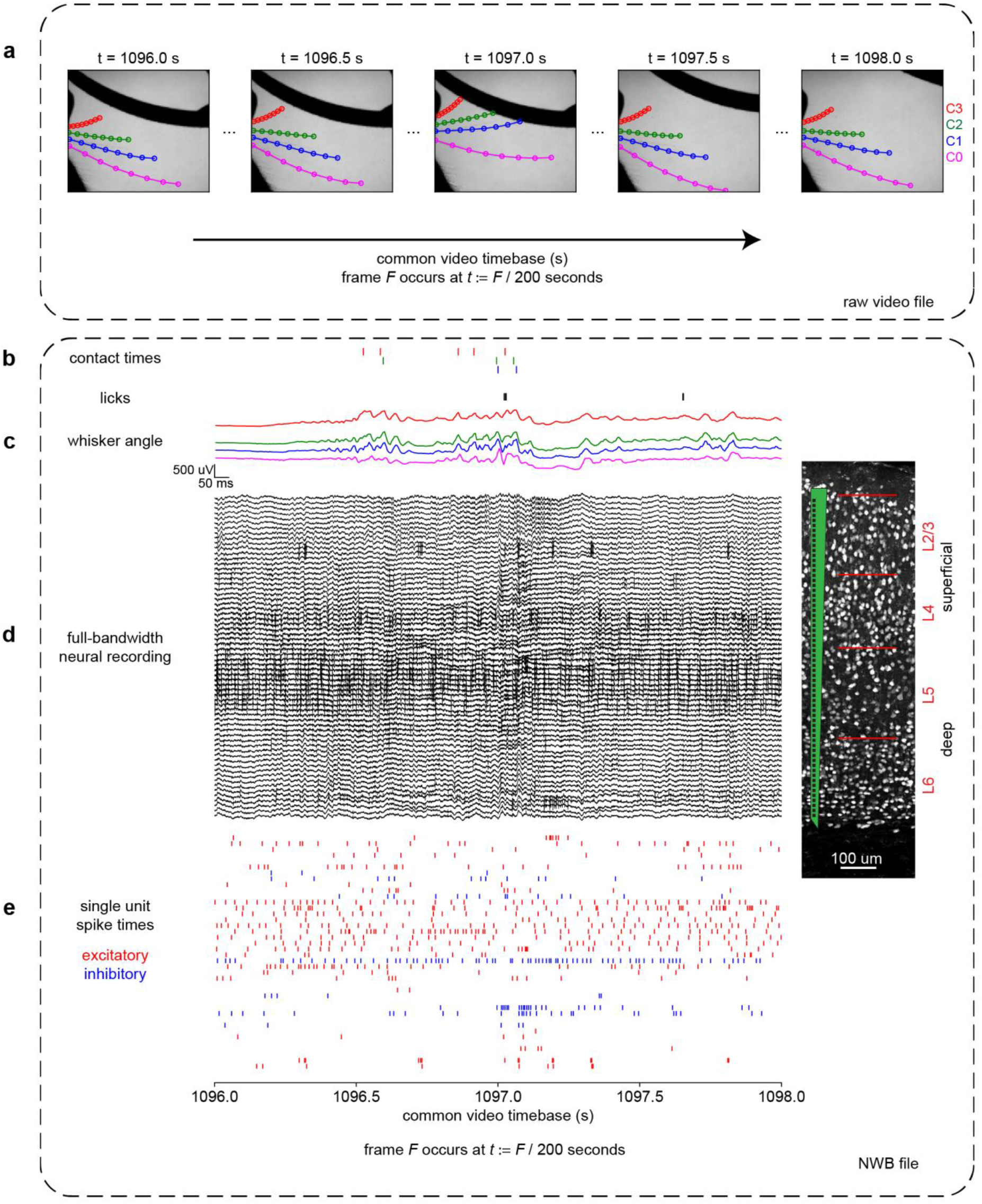
Description of dataset. Panel A shows data from the raw video whereas panels B-E show data from the NWB file. All panels show the same 2 seconds of aligned data from the same session. **A)** Typical frames within the video. 8 markers from each of 4 whiskers shown here, but not included in the raw video file. **B)** Times of whisker contacts (color raster) and licks (black raster). **C)** Angle of each of the four whiskers (colors as in panel A). **(D)** 64 channels of raw neural data. Highpassed at 3 Hz for display here. Spikes are visible as quick vertical deflections. *Inset*: A histological image adapted from Ref ^6^ with schematic of the 64-channel recording device. **E)** Spike times from identified individual excitatory (red) and inhibitory (blue) neurons.

This dataset will be useful for understanding the motor control strategies mice use to direct their whiskers toward objects of interest, and the way they interpret the resulting sensory input to make decisions. The inclusion of two different tasks permits an analysis of how the cortex can be reconfigured depending on task needs. Finally, our rigorously validated whisker tracking data will enable the development and testing of pose tracking algorithms, and our neural responses can be used to enable the development and testing of spike sorting algorithms. Thus, the data we present here can be used to open many avenues of investigations, distinct from the analyses we initially presented.

## Methods

### Two behavioral tasks

We trained two groups of mice to perform one of two tasks: shape discrimination (concave vs convex) or shape detection (either of those shapes, vs nothing). Mice were placed on a regulated water schedule to motivate them to perform the task to obtain water rewards. For each session, they were head-fixed and trained to lick either a left lickpipe or a right lickpipe in front of their face, depending on the presented stimulus. Shape discrimination mice were trained to lick left for concave shapes and right for convex shapes. Shape detection mice were trained to lick right for either shape (concave or convex) and to lick left on catch trials when no shape was presented at all. No mice were trained on both tasks.

On each trial, a linear servo motor moved a curved shape into the range of the whiskers on the right side of the mouse’s face. The shape stopped at one of three different positions (close, medium, or far). Even at the closest position, mice always had to actively move their whiskers in order to touch the shape. Mice performed this task in the dark, and we ensured that they relied on whisker-mediated touch to perform the task.

Mice could move their whiskers, make contacts on the shape, and lick left or right at any time during the trial. We took as their “choice lick” the first lick that occurred in a period of time called the “response window”, which began exactly 2 seconds after the linear servo motor started moving the shape toward the mouse. If we define t = 0 as the opening of the response window, then the shape begins moving toward the mouse at t = -2 s, and reaches its final position between -0.8 s and -0.4 s, depending on whether the final position was close, medium, or far.

Although we monitored all licks that mice made, all licks before the response window were irrelevant to the outcome of the trial. Only the first lick (left or right) during the response window determined the trial outcome (correct or incorrect). Correct licks were rewarded with water and incorrect licks were punished with a timeout, typically 9 seconds long.

Mice have five rows of whiskers, which are overlapping and difficult to distinguish in video. To permit unambiguous identification of whisker identity, we trimmed off all whiskers except the middle row, called the “C row” of whiskers. Within this row of whiskers, the longest and most posterior whisker is called C1, whereas the shortest and most anterior whisker still capable of reaching the shapes is called C3. We also tracked a longer and more posterior “straddler” whisker (beta or gamma, but referred to here as C0 for simplicity), but this whisker rarely made contact with the shapes in most mice.

The two tasks (discrimination and detection) were designed to be as similar as possible, other than the task rule. For each task, we used the same behavioral equipment, shape stimuli, trial timing, behavioral shaping, and whisker trimming procedures. The only difference between the tasks was the rule governing the correct response given the presented stimulus.

As described in our previous publication^6^, the original experiments were conducted at Columbia University, under the supervision and approval of the Columbia University Institutional Animal Care and Use Committee. We purchased breeder mice from the C57BL/6J strain (Jackson Laboratories Stock #000664), but all mice reported here were bred in-house. Mice began behavioral training between postnatal days 90 and 180. They were kept on a 12 hour, non-reversed light cycle and were typically tested during the day. They were typically group-housed with littermates until neural recordings began, and then they were singly-housed to protect the implant. They were provided with a running saucer (Bio-Serv InnoDome) for enrichment. Mice were assigned arbitrarily to shape discrimination or shape detection tasks. We used males and females arbitrarily and in roughly equal proportion. Female mice typically weighed less than male mice and drank correspondingly less water, but we adjusted the reward size based on weight to achieve roughly equal trial counts. Because we observed no other differences, we pooled the data from both sexes. During spike sorting and pose tracking, we were blinded to the trial outcomes, though not to task type or mouse identity.

### Session control, and trial types to be dropped from analysis

For both tasks, the behavior was controlled by a desktop computer connected to an Arduino. The computer ran custom Python software to choose which stimulus to present, and sent this information to the Arduino. The Arduino controlled the motion of the motors to present the correct stimulus, and delivered water rewards by opening a solenoid upon correct licks. The Arduino also reported all licks and rewards in plain text over a USB connection.

On most trials, the stimuli were chosen at random. These “random trials” are the primary data that we analyzed. However, on other “non-random trials”, the stimuli were chosen specifically to counteract undesired behavioral strategies and to encourage the mouse to do better. For instance, if the mouse repeatedly licked in the same direction (e.g., left) on trial after trial, then we would switch to non-random trials that required a lick in the opposite direction (right). In addition, if the mouse repeatedly licked in the same direction as on the previous trial (a so-called “stay bias” or perseverative bias), then we would switch to non-random trials that required the mouse to lick in the opposite direction as the last reward. We called this last mode of operation “Forced Alternation” because the optimal strategy is to alternate lick directions from trial to trial. Even in fully trained mice, we found it best to begin every session with 45 trials of Forced Alternation, before switching to random trials. All of these trials are included in the dataset we present here, but we have distinguished the “random trials” from “non-random trials”, and we suggest that only the random trials should be analyzed in most cases.

On some sessions, we also delivered optogenetic stimulation (laser light) to the brain for ongoing and unpublished optogenetics experiments. During these sessions, the laser was activated on a random subset of trials, and the remaining trials were normal. We identify these trials as “optogenetic” in the dataset, and we suggest that they should be dropped from analysis. This is described in more detail in the section “Data Records” / “Annotated data in the NWB file” / “NWB file: trials table”.

### High-speed videography and whisker tracking

In order to determine how mice explored the shapes and what sensory evidence they used to make their decision, we used a Photonfocus DR1-D1312IE-100-G2-8 camera to collect high-speed video of the whiskers as they interacted with the shape. All of the data in this dataset was collected after mice had learned to perform the task well with a single row of whiskers, not during learning. The camera captured 200 frames per second, and in most sessions the resolution was 640x550.

Using this video, our goal was to assess the entire extent of each of the four whiskers (C0-C3) on every frame, identify which whisker was which, and detect every contact made on the shape. The size of the dataset—15 mice, 88.9 hours, 115 sessions, 18,514 trials, 63,979,800 frames—required the used of high-throughput video analysis.

Most modern algorithms for tracking animal position require a training set, in which a human observer identifies the location of each discrete joint (such as a wrist) on a set of representative images. The animal’s rigid skeleton may then be inferred as the connections between those joints. Whiskers are more difficult to track because they are flexible and continuous. They lack any obvious anatomical features along their extent that can serve as discrete markers, other than the base and the tip. To address this, we used a previous-generation (non-neural network) whisker tracking algorithm called `whisk` to trace whiskers in individual frames^11^. `whisk` required no training data and it readily tracked the entire extent of all visible whiskers, though in our conditions it proved to be inaccurate at identifying which whisker was which. Therefore, a human observer (CR) curated its output by discarding poorly tracked or spurious whiskers (such as fur), and by labeling each whisker C0-C3. Then, we extracted 8 equally spaced markers along these curated whiskers (Fig 1A). These markers do not represent any physical attribute of the whisker but are simply defined to be equally spaced. In this way we generated an accurate training set of 8 labeled markers for each of 4 whiskers, suitable for training a modern pose tracking neural network.

We used a deep convolutional neural network developed for tracking the position of human body parts^12,13^, and also shown to be effective for tracking animal pose^14^. After training the network on our curated training set, we used it to predict the whisker positions on the entire unlabeled videos. While the network was accurate on most frames, it occasionally made large errors when the animal was whisking intensely, which were exactly the frames of greatest interest. These types of rare but severe errors are obscured by a standard “RMS pixel error” metric, and for that reason we favor a “frame-wise error probability” metric (i.e., the fraction of frames on which the error exceeded some predefined threshold of acceptability).

To address this, we iteratively 1) selected unlabeled “challenge frames” that were difficult to predict; 2) corrected these “challenge frames” by choosing the output of either the network or `whisk` and by labeling the whiskers manually; and 3) retrained the network with the corrected challenge frames added to the previous training set (Figure 2). We repeated this process 5 times until the whisker-wise error probability reached <0.2%, assessed using doubly held-out cross-validation (see “Validation of pose tracking” in “Technical Validation”).

**Figure 2.**
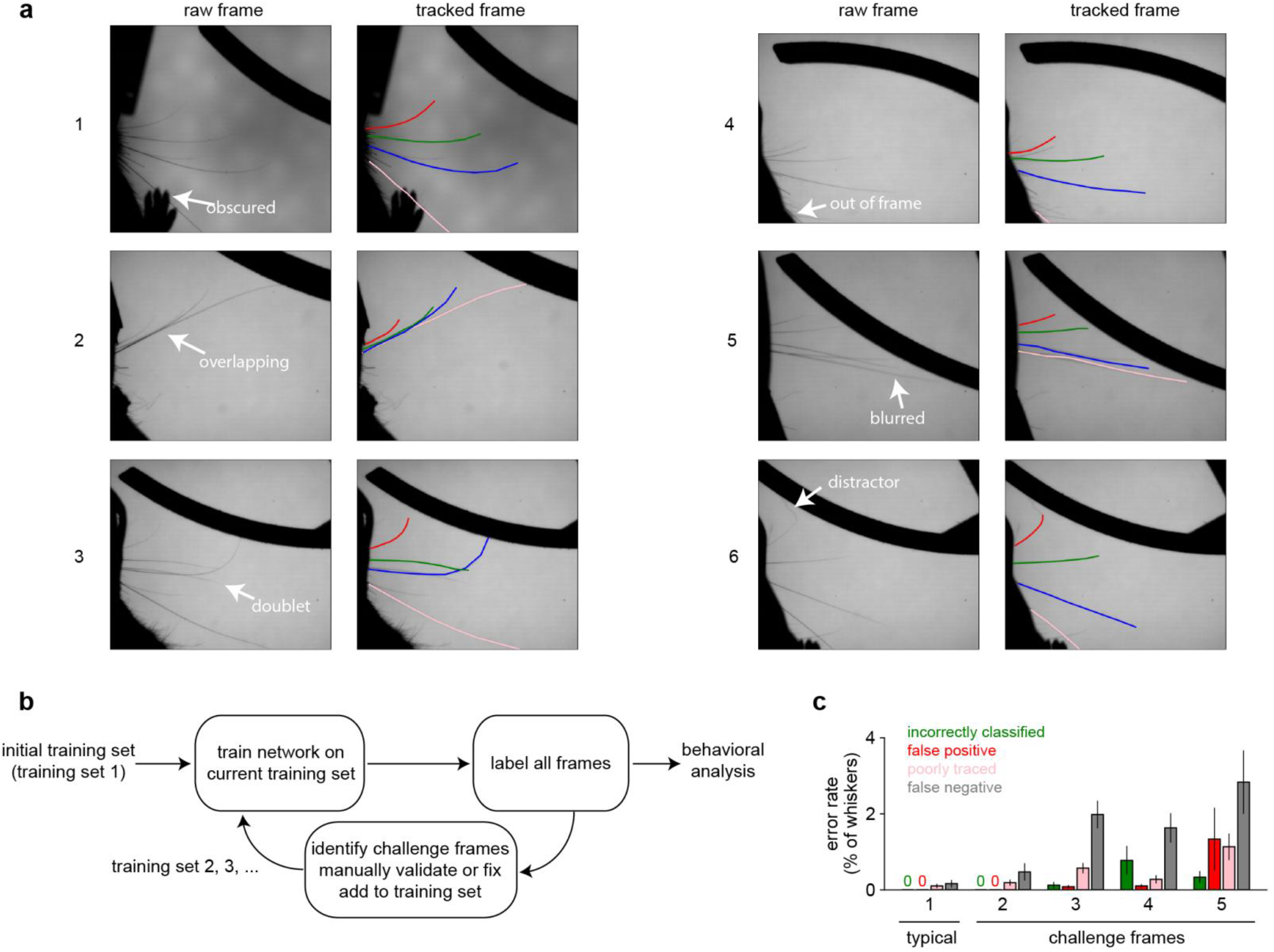
An iterative method for training a pose tracking network. This figure is adapted from Supplemental Figure 2 in Ref^6^. **A)** Example frames demonstrating the quality of the whisker tracking. Within each pair of frames, the left frame is the raw frame (annotated with the region of interest) and the right frame shows the result of the whisker tracking algorithm. Performance was good (i.e., the correct whiskers were tracked throughout their extent) even when: 1) the whisker was obscured by a paw; 2) whiskers were nearly overlapping; 3) a “doublet” whisker emerged from the same follicle; 4) the whisker was nearly out of frame; 5) motion blurred the tips; 6) a similar-looking distractor hair was attached to the end of the whisker. **B)** We used an iterative procedure to train the network, repeatedly identifying “challenge frames” on which it struggled and using those to generate the next training set. **C)** Error rates for the initial training set of typical frames (training set 1), as well as each set of increasingly difficult challenge frames (training set 2-5).

Finally, we used the tracked extent of the whisker to estimate its bending (i.e., spatial curvature) as it contacted objects, a commonly used proxy for contact force. To do this, we fit a third-order spline through the eight markers and passed the spline to the `measure` function of `whisk`. The end result of this process was a detailed and accurate picture of the position of all four individually tracked whiskers, even during periods of rapid motion, spatial overlap, or intense contact with the object.

### Neural recordings during behavior

For a subset (44/115) of the behavioral sessions we report here, we also recorded neuronal activity in identified columns of the barrel cortex. Located within the primary somatosensory cortex, the barrel cortex is well-known for its somatotopic organization, meaning that each whisker on the face corresponds topographically to a specific column in the barrel cortex. Decades of research have studied how computations in this area contribute to tactile perception and decision-making^15^. However, few studies have combined recordings with high-speed video of multiple identified whiskers during a goal-directed behavior.

Briefly, once mice learned the task, we anesthetized them and located the left barrel cortex through the thinned skull. Specifically, we stimulated individual whiskers using a piezo and monitored the reflectance of the brain to red light (an indicator of deoxygenated blood flow and hence neural activity) to identify the specific cortical column showing the earliest and strongest response to the whisker stimulation. Thus identified, we aligned the cortical columns corresponding to each individual whisker to a picture of the vasculature map to guide future recordings. Next, we cut a small craniotomy over the barrel cortex (typically including columns C1, C2, and C3) using aseptic technique. We sealed the craniotomy with silicone gel (Dow DOWSIL 3-4680, sometimes called “artificial dura”) and a silicone sealant (WPI KwikCast). We also implanted a stainless steel pin elsewhere on the dorsal surface of the skull, making electrical contact with the brain without actually penetrating it, to serve as a combination ground and reference. Once the mouse recovered from surgery, we resumed the behavioral experiment, now concurrent with neural recording.

During each recording session, we head-fixed the mouse in the behavioral arena as usual, and then removed the silicone sealant, thus exposing the identified cortical columns. We used a micromanipulator to position the electrode perpendicular to the cortical surface, noting its location within the barrel cortex using the vasculature map. We slowly lowered the electrode into the brain, noting its depth and final location within the cortex. Once positioned, we began acquiring neural data with an Open Ephys recording system^16^, with the headstage filters set to the maximum possible bandwidth (1 Hz to 7.5 kHz). After the behavioral session, we ended the recording, carefully raised the electrode out of the brain, and sealed the craniotomy again as before. In this way, during each behavioral session we knew the precise location of the electrode within the cortical map, and also its precise depth within the cortex. On subsequent recording sessions, we would sample a different cortical column, with the goal of sampling columns C1, C2, and C3 equally often.

We used a silicon probe (Cambridge Neurotech H3) with 64 extracellular recording sites arranged linearly from superficial to deep (Figure 1d). The distance between site centers was 20 μm and the site area was 11 μm by 15 μm, coated with PEDOT to a target impedance of 50 kΩ by the manufacturer. These design parameters permitted us to isolate individual neurons (“single units”) based on their extracellular waveform shape over multiple recording sites. The 64 recording sites spanned 1280 μm of cortical depth, allowing us to simultaneously sample almost the entire thickness of this cortical region, from the superficial layer 2/3 (L2/3) to the deep layer 6 (L6). We defined the layer boundaries with published measurements^17^.

To isolate individual neurons (“spike sorting”), we used the software Kilosort^18^, which uses a template matching approach that is suitable for high-density electrode arrays, especially in the case of spatially and temporally overlapping spikes. We used the default parameters, except for changing the number of templates to 192 (*i*.*e*., thrice the number of channels). We used the software Phy^19^ to manually inspect the results of Kilosort. Using the waveform geometry and cross-correlograms, we identified cases where the same neuron had been artefactually split into two different templates (most frequently because of slight spatial drift along the electrode array over the session) and merged such templates. Using the waveform geometry and auto-correlograms, we preliminarily identified “well-sorted” units as opposed to multi-unit templates (recognizable by an auto-correlogram peaking near time zero) and noise templates (recognizable by a large and similar waveform across all channels of the array). These subjectively-identified “well-sorted” units were then subjected to strict and objective quality tests, described in “Technical Validation” / “Spike Sorting”.

### Ordering of neural data channels

The Open Ephys acquisition system stores the channels of neural data in the order defined by the pin numbers on the Intan headstage, which bears no relation to the location of the recording sites in the brain. Moreover, during most recording sessions we identified at least one “broken channel” that produced no useful data, typically because of a flaw in the recording site or headstage. Finally, the depth of the array within the brain could differ from day to day, so the same channel did not always record from the same depth within the brain.

Therefore, for ease of use, we ordered the channels in the Neurodata Without Borders (NWB) file in order of the anatomical location of the corresponding recording site, from superficial to deepest. We excluded broken channels from the NWB file. The included channels are specifically identified in the `electrodes` table of the NWB file, as defined further below. The neural data in the `acquisition` group is stored in exactly the same order as the rows of the `electrodes` table. The location of each individual neuron is defined by a field called “neuron_channel” in the `units` table, which is an index into the `electrodes` table identifying the channel on which that neuron was recorded. Finally, the `electrodes` table contains a string called “location”, which identifies for each channel its anatomical location (depth within cortex) as well as the Intan headstage channel number. Including this Intan channel number allows users to go back to the raw data in the Open Ephys files and find the same channel of neural data.

### Alignment of multiple data streams

A strength of this dataset is that it combines behavioral data (*e*.*g*., stimulus and choice times), video data (whisker positions and contacts), and neural data (spikes and local field potential). However, these distinct data streams must be properly and carefully aligned to permit analysis. Each data stream has its own clock. The behavioral data is recorded in milliseconds since the beginning of the behavioral session according to the Arduino. The video data is recorded in frames (at 200 frames per second) since the beginning of the video session. The neural data is recorded at 30 kHz since the beginning of the neural recording. The behavioral, video, and neural sessions did not begin at exactly the same time, and the clocks on these various devices do not run at exactly the same speed. Finally, we observed that the Arduino’s clock in particular varied its speed over the course of the session, perhaps due to fluctuating temperature levels.

To account for these challenges, we used a single synchronization pulse that occurred at the beginning of every trial. The Arduino that controlled the behavioral task set low one of its normally high digital outputs for 133 ms at the beginning of each trial, and noted the time of this in the behavioral logs. This digital output was directly sampled by an analog input in the Open Ephys acquisition system, and it was also connected to a solid-state relay (SSR) that switched off the backlight underneath the mouse whiskers. Thus, the backlight turned off for 133 ms at the beginning of each trial, causing the video data to go completely black for that time.

This method created a single, unambiguous signal at the exact start time of each trial that could be observed in the behavioral, neural, and video data. Specifically, we identified 1) the start time of each trial in the behavioral logs, 2) the exact sample at which the correspond analog input went low in the neural recording, and 3) the exact frame on which the backlight went dark on each trial. The variable time between trial starts provided a unique cue to which trial was which. In this way we could identify the same moment at the start of each trial in each data stream (Figure 3).

**Figure 3.**
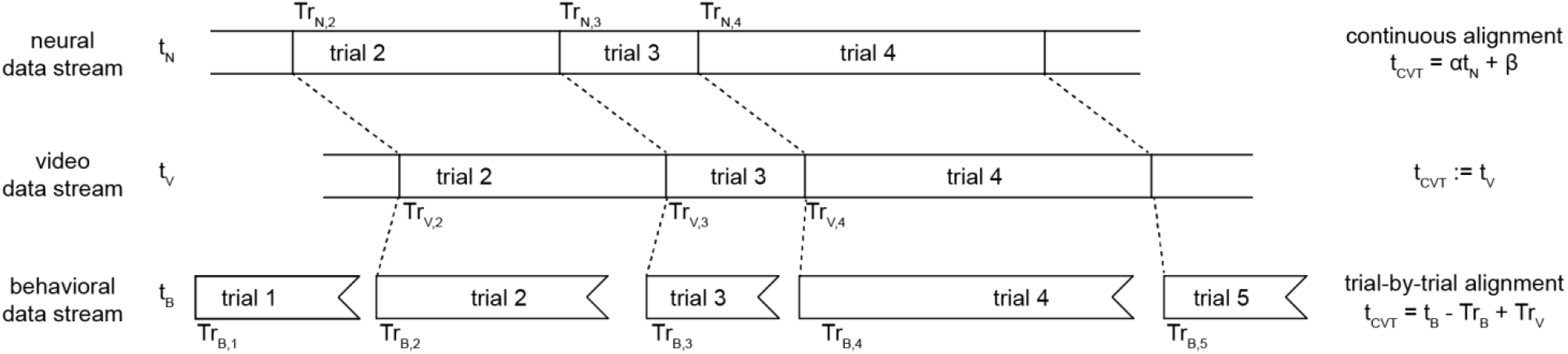
Schematic of procedure for aligning data streams This dataset contains three streams of raw data: neural recordings, video frames, and behavioral events. They are aligned by a synchronization pulse at the beginning of each trial, which is detected in all three streams at times *Tr*_*N,i*_, *Tr*_*V,i*_, and *Tr*_*B,i*_ for trial *i*. The time between trials varies, allowing the same trial to be matched up across all streams, although the neural and video streams may be missing trials at the beginning or end of the session. To align the neural timebase (top) to the common video timebase (middle), we used a continuous alignment procedure. To align the behavioral timebase (bottom) to the common video timebase (middle), we had to use a trial-by-trial alignment procedure, which introduces slight discontinuities (no more than a few milliseconds) between trials.

Throughout this dataset, we always aligned behavioral and neural data to the video data, rather than the other way around. That is, we mapped the time of behavioral events and neural events to our best estimate of when each event occurred in the video. We refer to this time in the video as the “common video timebase” throughout this manuscript. Since the camera ran at 200 frames per second, frame F in the video is defined as occurring at exactly time F/200 in the common video timebase.

However, this alignment can be performed in two different ways. The first way is trial-by-trial: the time of every event is normalized by subtracting the most recent trial start time, and then this normalized value is added to the trial start time in the common video timebase (t_CVT_ = t_X_ – T_start,X_ + T_start,CVT_). The second way is continuous: a linear fit is obtained between the trial start times in two different timebases (*e*.*g*., neural and video), and then this linear fit is used to smoothly transform times between the two (t_CVT_ = αt_X_ + β).

The trial-by-trial method is more robust to gradual changes in the relative clock rates across the session, such as we observed in the behavioral timebase due to the Arduino, but it also creates small discontinuities at the boundaries between trials. Also, the trial start times can only be estimated to the nearest frame (5 ms) in the common video timebase, which limits the precision of this method.

In contrast, the continuous method can achieve sub-frame precision by fitting the same linear equation to all trial start times. However, this will only be accurate if the two timebases are consistently related over the entire session—an assumption we know to be false for the Arduino, but which was tenable for the video and neural clocks (see “Technical Validation” / “Alignment data”). Therefore, we used the trial-by-trial method to align the behavioral data to the common video timebase, and the continuous method to align the neural data to the common video timebase.

## Data Records

The data we present here consists of three general categories: behavioral events, video of the whiskers, and neural recordings. A major technical challenge in analyzing these data is synchronizing across distinct data streams. For instance, a user may wish to identify all trials with concave shapes from the behavior data, and slice out aligned video and neural recordings from each of those trials. To enable easier analysis of these raw data, we have generated and carefully annotated a single Neurodata Without Borders (NWB) file for each session that combines synchronized and processed versions of the behavioral, neural, and video data for that session.

We used the Dandi Archive to store the data because it provides a standardized interface for users to download part or all of the dataset^9^. Users with limited bandwidth do not have to download any data to get started: they can run analyses on Dandi Hub, which includes browser-based widgets for exploring and analyzing the data, such as viewing spike trains or plotting PSTHs to whisker contacts or trial-based events. The entire dataset is available at https://dandiarchive.org/dandiset/000231.

### Directory structure of the entire dataset

Dandi uses a standardized interface for naming and structuring data directories. At the top level, there is one folder per subject (*i*.*e*., mouse) named “sub-mousename”, where “mousename” is a unique name for that mouse. Each subject’s folder contains multiple NWB files, one per session from that mouse. Each NWB file is named “sub-mousename_ses-sessionname_behavior+image.nwb”, where “sessionname” is a unique name for that session. The string “behavior+image” indicates that all sessions include behavior data and whisker video.

Some sessions also include extracellular electrophysiology data, in which case the filename includes “ecephys” as well.

For technical reasons, NWB files are not suitable for storing behavioral videos such as ours. Therefore each NWB file contains a link to an external raw video file, one per session. Video files are stored in a directory with the same name as the corresponding NWB file.

Finally, we also provide the raw neural data files in a directory called “sourcedata” whose subdirectories use the same “sub-mousename” structure defined above. The neural data include Open Ephys data files, exactly as we acquired them, and also the results of spike sorting with Kilosort and Phy.

In the following sections, we describe the components of the NWB file, which we believe will be the most useful for most users. We then briefly describe the raw video and neural data files.

### Annotated data in the NWB file

The NWB file format is a standard format for exchanging neuroscience data^9^. It prescribes specific ways to store data, to encourage best practices (such as storing all metadata alongside the data itself) and to make it easier for users familiar with one NWB dataset to use another one. The NWB file itself stores data in HDF5, which is hierarchical (tree-like). Some fields are simple items of metadata (*i*.*e*., start time of the session) and other fields are groups, potentially deeply nested (*i*.*e*., an acquisition group containing all of the acquired neural data).

At the top level, the NWB file comprises several pre-defined fields. This is documented by the maintainers of NWB^9^. In Table 1, we identify only the fields that require further explanation in our dataset. We provide that further explanation in each sub-section below.

**Table 1.** List of field names and descriptions at the top level of the NWB file.

| Field | Description |
| --- | --- |
| acquisition | A group containing `ElectricalSeries` objects for neural recordings, and `ImageSeries` objects for behavioral videos. |
| electrodes | A `DynamicTable` specifying characteristics of each electrode |
| processing | A group containing a `ProcessingModule` named “behavior”. This includes processed data about behavioral events, whisker position, and whisker contacts. |
| trials | A `TimeIntervals` specifying the parameters of each of the behavioral trials in this session |
| units | A `Units` specifying the recorded neurons and their spike times |

#### NWB file: acquisition group: “extracellular array recording”

Raw neural recordings are contained in a single `ElectricalSeries` named “extracellular array recording”.

This `ElectricalSeries` contains the following fields:

**Table 2.**
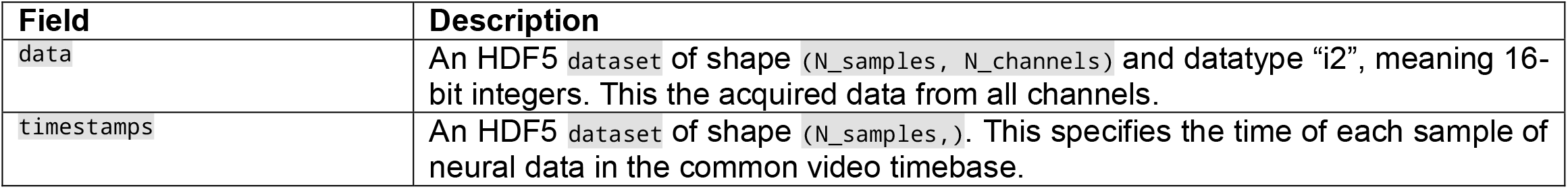
List of field names and descriptions within the `acquisition` group “extracellular array recording”.

The section “Temporal alignment of all data streams” describes how the timestamps were calculated. The section “Ordering of neural data channels” describes the order of the channels of data with respect to the recording sites in the brain.

The raw neural data may be accessed as follows:

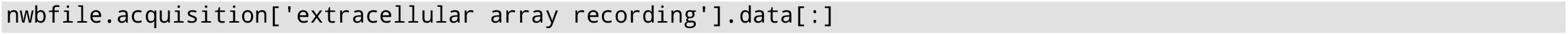

This returns a numpy array of continuous neural data from all channels and all timepoints.

The timestamps may be accessed as follows:

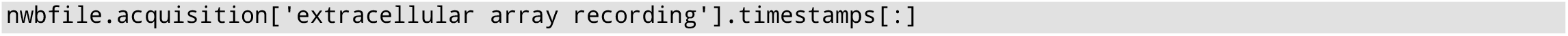

This returns a numpy array of regularly spaced timestamps, one per sample of neural data. The sampling rate is nearly 30 kHz, but not exactly, because it has been converted to the common video timebase used throughout the NWB file.

#### NWB file: acquisition group: “behavioral video”

This `ImageSeries` represents the acquired behavioral video of the whiskers. Because it is not feasible to store video in an NWB file, this simply includes the relative path to the video file.

The filename may be accessed as follows:

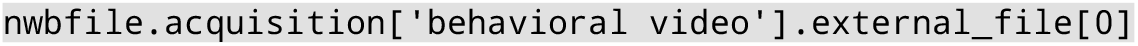

#### NWB file: processing group: “behavioral_events”

This `ProcessingModule` includes two data interfaces of type `BehavioralEvents`. The data interface named “licks” comprises two `TimeSeries`, one named “left” and one named “right”, specifying the times of left and right licks. (On rare occasions when mice licked both lickpipes at the same time, this is stored in a third `TimeSeries` called “both”.) Similarly, the data interface named “rewards” comprises two `TimeSeries` named “left” and “right”, specifying the times of left and right rewards.

The times of licks and rewards may be accessed as follows:

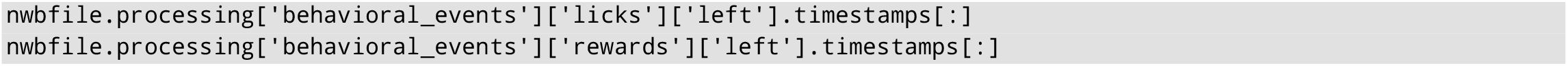

These return numpy arrays of lick or reward times, expressed as the number of seconds in the common video timebase.

#### NWB file: processing group: “raw_whisker_position”

This `ProcessingModule` contains the location of tracked body parts estimated by the pose tracking network. It comprises four data interfaces, each of which is an `ndx_pose.pose.PoseEstimation` containing pose tracking data for one of the four whiskers that we tracked. In turn, each `PoseEstimation` comprises eight `PoseEstimationSeries`, one for each of the eight markers that we tracked along the length of that whisker. Low-confidence markers are excluded, and so there are frames for which some or all data is missing.

The location of marker 7 along whisker C2 (for example) may be accessed as follows:

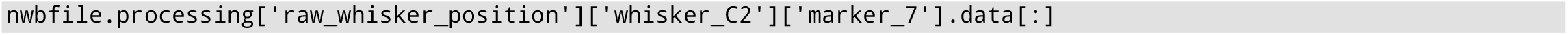

This returns a numpy array of shape `(N_frames, 2)`. The first column gives the row index of the marker in pixels and the second column gives the column index of the marker in pixels. These pixel values may be converted to millimeters by multiplying by the value in the `conversion` field of each `PoseEstimationSeries`, which is the same for all markers within a session but can vary across sessions.

#### NWB file: processing group: “processed_whisker_position”

This `ProcessingModule` contains information about the location of each whisker after processing the data in the group named “raw_whisker_position”. Specifically, we interpolated the whisker positions over frames in which the data were missing because the pose tracking network’s confidence was too low to make a prediction. Also, we calculated the angle and the bend of each tracked whisker. The bend is a measure of curvature, and increases as the mouse pushes its whisker against a shape.

This group comprises four data interfaces, each of which is a `BehavioralTimeSeries` containing data about one of the four tracked whiskers on each frame. In turn, each `BehavioralTimeSeries` contains various `TimeSeries` of the same length.

**Table 3.** Names and descriptions of the `BehavioralTimeSeries` within each whisker’s data interface in the processing group “processed_whisker_position”.

| Name (Type) | Description |
| --- | --- |
| angle (float) | The angle between base and tip of the whisker, with 0 meaning horizontal and positive values indicating protraction |
| base_x (float) | The x-coordinate of the base of the whisker, in pixels |
| base_y (float) | The y-coordinate of the base of the whisker, in pixels |
| bend (float) | The bend of the whisker (units: $m^{-1}$), with 0 meaning straight and positive meaning pushing into an object. This is null on interpolated frames. |
| interpolated (bool) | A bool indicating whether this frame was interpolated over missing frames or not |
| tip_x (float) | The x-coordinate of the tip of the whisker, in pixels |
| tip_y (float) | The y-coordinate of the tip of the whisker, in pixels |

The angle of whisker C2 (for example) may be accessed as follows:

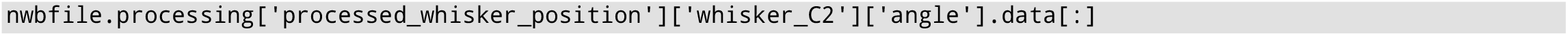

This returns a numpy array of shape `(N_samples,)`, where `N_samples` is the number of frames with data after interpolating over missing frames.

The timestamps may be accessed as follows:

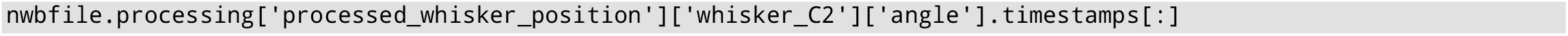

This returns a numpy array with the same shape as the data in the `BehavioralTimeSeries`, specifying the time of each data point in the common video timebase. Rather than covering the entire session evenly, these timestamps will in general contain gaps, for instance during the 133 ms blackout synchronization pulse at the start of each trial. This is because we did not interpolate over gaps longer than 4 frames (20 ms).

#### NWB file: processing group: “identified_whisker_contacts”

This `ProcessingModule` contains information about the time, location, and kinematic parameters of every contact made by each whisker on the shape stimuli. Because the mouse chose when to make contacts by actively whisking, contacts could occur at any time, and any given trial could contain any number of contact events. Therefore we used a `TimeIntervals` object to specify the timing of each contact event.

This group comprises four data interfaces, each of which is a `TimeIntervals` containing data about contacts made by one of the four tracked whiskers. The required fields “start_time” and “stop_time” specify the beginning and end of the whisker contact. We have also added optional fields containing the location of each contact, expressed both as a Cartesian location within the frame and also as the angle of the whisker at the time of the contact.

These data may be accessed as follows:

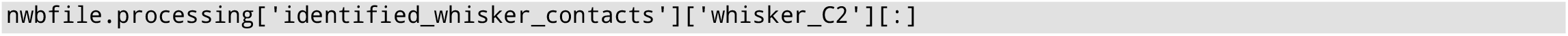

This returns a pandas DataFrame with shape `(N_contacts, N_fields)`, where `N_contacts` is the number of contacts made by that whisker and `N_fields` is the number of fields of data about each contact (start time, stop time, *etc*.).

#### NWB file: electrodes table

This is a `DynamicTable` specifying the metadata about each electrode (*i*.*e*., channel on the silicon array). The fields in this table are prescribed by the NWB standard.

**Table 4.** Names and descriptions of the columns of the electrodes table.

| Name (Type) | Description |
| --- | --- |
| y (float) | The depth of the electrode in microns from the surface of the brain. The related fields “x” and “z” are not relevant for this recording device and are left as null. |
| location (string) | We use this string to encode the laminar location of the electrode and its channel number in the original data file. This is documented further below. |
| electrode_group<br>(`ElectrodeGroup`) | We use only one `ElectrodeGroup` in each session. Its “location” field specifies the barrel-related cortical column in which the recording device was inserted during this session (e.g., “C2”). |

The number and order of the electrodes in this table exactly matches the number and order of the channels of raw data in the acquisition group. Broken channels are excluded from both the acquisition group and the electrodes table. The field “neuron_channel” in the table `nwbfile.units` indexes this electrodes table to specify where each neuron is located. The section “Ordering of neural data channels” describes the order of the channels of data with respect to the recording sites in the brain.

Because the electrodes table cannot contain additional custom columns of information, we encoded additional metadata about each data channel in the string contained in the “location” field. This string includes in human readable text the following metadata:

- the laminar location in cortex (based on the depth),
- the channel number within the raw Open Ephys data files (based on the Intan headstage),
- the channel number in the Open Ephys GUI (which is always 1 greater than the Intan headstage),
- the channel number in anatomical order (superficial to deepest) including broken channels,
- and the channel number in anatomical order (superficial to deepest) excluding broken channels.

The difference between the last two indexes is whether broken channels are included. For example, if channel 2 was broken the first ordering would be [0, 1, 3, …, 63] and the second would be [0, 1, 2, …, 62]. The first method is advantageous for identifying the same physical channel across days, while the second method is advantageous for indexing the rows of the electrodes table without gaps.

These data may be accessed as follows:

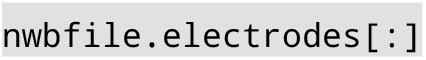

This returns a `pandas.DataFrame` of shape `(N_working_electrodes, N_fields)`, where `N_working_electrodes` is the number of working channels in this recording session (out of a maximum of 64) and `N_fields` is the number of fields of metadata about those channels.

#### NWB file: units table

This is a `Units` table that specifies metadata about each identified single neuron (“unit”) in the data.

**Table 5.**
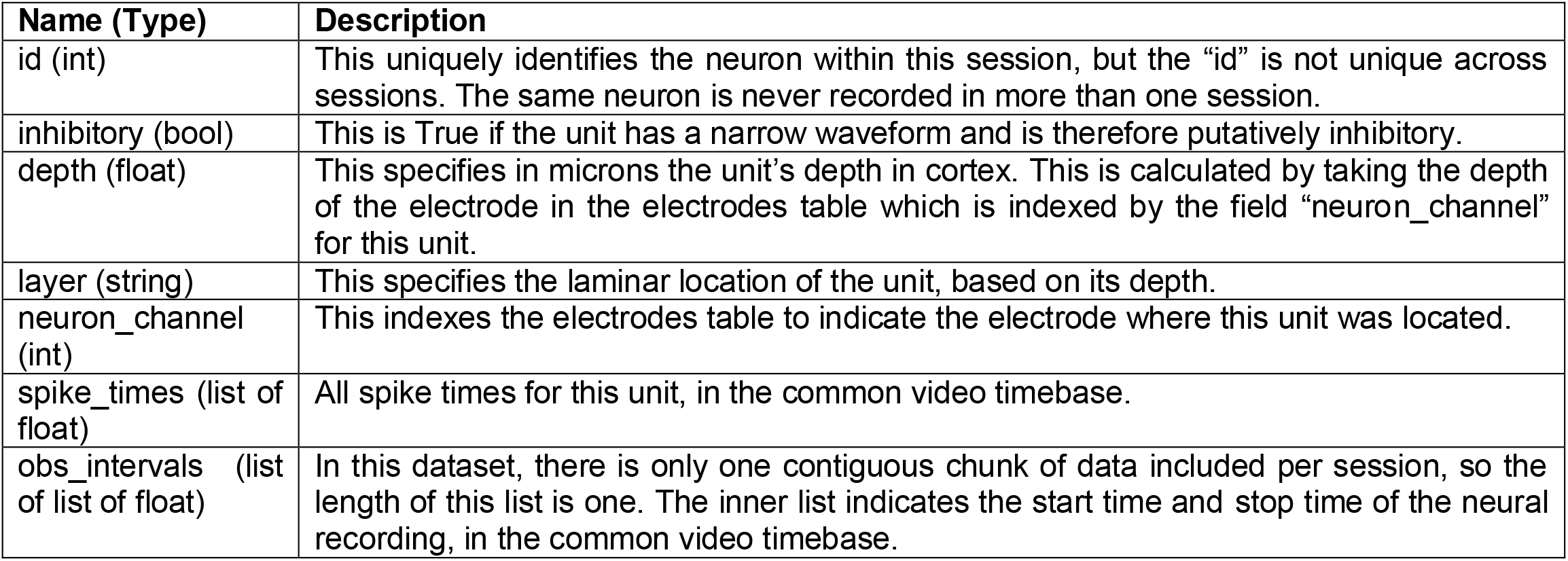
Names and descriptions of the columns of the units table.

We defined the location of each neuron as the channel on which that neuron’s waveform had maximal root mean square power. This electrode is specified by the field “neuron_channel” in this table, which indexes both the electrodes table and also the channels of raw data in the acquisition group. That is, if the field “neuron_channel” is 0 for a given unit, then the unit was located on the first electrode in the electrodes table, and that electrode’s recording is in the first column of data of the acquisition group.

The spike times were converted to the common video timebase by continuous alignment of the data streams, not by trial-by-trial alignment. This means that the spike times given here may be directly compared to the timestamps of the raw neural data in the acquisition group.

These data may be accessed as follow:

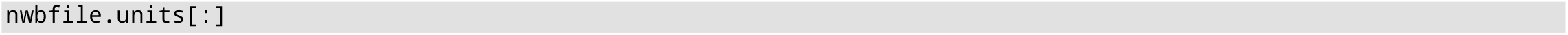

This returns a `pandas.DataFrame` of shape `(N_units, N_fields)`, where `N_units` is the number of identified single neurons in this recording session and `N_fields` is the number of fields of metadata about those units.

#### NWB file: trials table

This is a `TimeIntervals` specifying metadata about each behavioral trial. For each trial, we provide fields that are required by the NWB specification (“start_time” and “stop_time”) as well as several custom fields.

**Table 6.** Names and descriptions of the columns of the trials table.

| Name (Type) | Description |
| --- | --- |
| start_time (float) | Start time of the trial |
| stop_time (float) | Stop time of the trial. Equal to the start time of the next trial |
| direct_delivery (bool) | If True, water was delivered to the rewarded pipe regardless of the mouse’s choice. Such trials should be ignored. |
| stim_is_random (bool) | If False, the stimulus identity was not chosen randomly, for the purpose of behavioral shaping. Such trials should be ignored. |
| optogenetic (bool) | If True, a laser was used to manipulate neural activity. Such trials should be ignored. |
| outcome (string) | One of {‘correct’, ‘error’, ‘spoil’}. Spoiled trials are those on which no choice was made and should be ignored. |
| choice (string) | One of {'left', 'right', 'nogo'}. 'nogo' indicates that no choice was made and the trial was spoiled. |
| rewarded_side (string) | One of {'left', 'right'}. Specifies which direction the mouse should have licked in order to receive reward. |
| servo_position (string) | One of {'close', 'medium', 'far'}. Specifies the location of the shape with respect to the mouse's face. |
| stimulus (string) | One of {'convex', 'concave', 'nothing'}. 'nothing' was only used in the detection task, not in the discrimination task. |
| ignore_trial (bool) | This is True if 'direct_delivery' is True, or 'stim_is_random' is False, or outcome is 'spoil', or 'optogenetic' is True, or the trial was otherwise munged. Trials for which this is True should be ignored. |
| choice_time (float) | The time of the choice lick, which is the first lick after the response window opened. On spoiled trials, this value is 45 seconds after the response window opened. |
| response_window_open_time (float) | The time at which the response window opened. The first lick after the response window opened determines the outcome of the trial. Licks before the response window opened did not affect the trial. |
| trial (int) | The trial number. |

All times are specified in the common video timebase. They were aligned using a trial-by-trial alignment procedure, as described in the “Methods” section “Alignment of multiple data streams”.

Trials for which the field “ignore_trial” is True should be excluded from analysis, typically because they were used for behavioral shaping (such as non-random or direct delivery) or for optogenetic stimulation. Rarely, trials were “munged” (e.g., the motor failed to move the shape, or other experimental glitch), and on such trials the field “ignore_trial” is also set to True.

In some cases the video recording was not started until after the first trial. Trials before the video began are excluded from this trials table. Therefore the first trial number in this table is not necessarily 0.

These data may be accessed as follow:

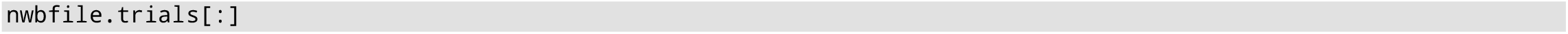

This returns a `pandas.DataFrame` of shape `(N_trials, N_fields)`, where `N_trials` is the number of trials in this session and `N_fields` is the number of fields of metadata about those trials. The index of this DataFrame is called “id” and is identical to the field “trial”.

### Raw data

#### Raw data: video files

For each session, we provide a video file that shows the whiskers and shape stimuli. The video data show how the mice moved their whiskers and interacted with the shape stimuli during the task. The video files can be played with any standard video player. We encoded them with the x264 codec into an mkv container using the command `ffmpeg -r 30 -pix_fmt yuv420p -vcodec libx264 -qp 15 -preset medium` and in most cases version 4.0.2 of `ffmpeg`. We chose the compression parameters (`-qp 15`, considered extremely high quality) such that the result was visually nearly indistinguishable from the original video.

Importantly, the frame rate of this video file is spuriously set to 30 frames per second, even though the true frame rate is 200 frames per second. That is, the video file plays at 30/200, or 0.15x, of the actual real-time speed. We did this because standard video players cannot reliably play back videos at high frame rates.

In the NWB file, all timestamps are specified in the common video timebase. This timebase is defined as a frame rate of exactly 200 frames per second. Thus, frame N in this video file occurs at exactly t = N / 200 seconds in the common video timebase, by definition.

To work with raw video files, we recommend using `ffmpeg-python`. This module allows extracting individual frames or chunks of frames for further processing, and is implemented using `ffmpeg`. However, any standard video reader should be capable of loading the video.

#### Raw data: neural acquisition files

The neural data show how the barrel cortex responded during the behavioral sessions. We only recorded neural activity in a subset of the sessions included in this dataset. The data were acquired by an Open Ephys acquisition system and stored directly in an Open Ephys file format (version 0.4).

These data are stored in the neural timebase, which is defined by Open Ephys as the number of seconds since it began streaming (but not necessarily saving) data. These data were stored in their entirety, including broken or disconnected channels, and without applying any filtering or rearranging of channels. In some cases the Open Ephys file includes “test recordings” that were just a few seconds long. The main recording was not guaranteed to begin with an particular alignment to the video or behavior, and may in fact have begun or ended too early or too late.

We have added all of the relevant neural data to the NWB file. The NWB file synchronizes the neural data to the common video timebase, excludes broken channels, excludes epochs of neural data that should not be analyzed (*e*.*g*., test recordings), and includes only well-sorted single units. Therefore we believe most users will rely on the NWB file, not the Open Ephys files, but we provide them for completeness.

## Technical Validation

### Spike sorting

As reported previously^6^, we used KiloSort^18^ to detect spikes and to assign them to putative single units. Single units had to pass both subjective and objective quality checks. First, we used Phy^19^ to manually inspect every unit, merging units that appeared to be from the same origin based on their amplitude over time and their auto- and cross-correlations. Units that did not show a refractory period (*i*.*e*. a complete or partial dip in the auto-correlation within 3 ms) were deemed multi-unit and discarded. Second, single units had to pass all of the following objective criteria: ≤5% of the inter-spike intervals less than 3 ms; ≤1.5% change per minute in spike amplitude; ≤20% of the recording at <5% of the mean firing rate; ≤15% of the spike amplitude distribution below the detection threshold; ≤3% of the spike amplitudes below 10 μV; ≤5% of the spikes overlapping with common-mode artefacts.

### Validation of pose tracking data

As reported previously^6^, we rigorously validated the accuracy of our whisker tracking, because we sought to have no discontinuities in tracking or missing contact events. The simplest metric of tracking accuracy is the mean distance between the true label and the predicted label, which is 2.8 pixels in our case. However, this is not a particularly informative metric, because the scientific utility of such an algorithm is limited by the prevalence of rare but large errors (e.g., misclassifying C1 as C2) which are too rare to affect such a metric. Also, there are multiple additional kinds of errors to consider, such as false negatives and false positives, each of which is more or less problematic depending on the analysis.

Moreover, performance on the *typical* frame is not particularly important: in the majority of frames in the dataset the whiskers are at rest or near rest, and in those frames tracking is easy. More rarely, mice would intensely whisk and touch the shapes in diverse ways, and although this constitutes a minority of the frames in the dataset, these are precisely the frames for which accurate tracking is most difficult and most important. In a catch-22, while frames with intense whisking and diverse touch dynamics are crucial for training the pose tracking algorithm and evaluating its performance, it is not obvious how to identify such frames without a working pose tracking algorithm. Thus, for all these reasons, quantifying the accuracy of tracking algorithms in general remains challenging.

We began by defining four disjoint types of whisker tracking errors, all defined with respect to a training set curated by a human (CR).

- “poorly traced”: The extent of the whisker is not traced correctly. For example, the tip is missing, or the trace “jumps” from one whisker to another. This was defined by calculating the Cartesian distance between ground truth and reported location of each joint in the whisker, and identifying whiskers where this distance was greater than 20 pixels for the tip or for the mean over all joints.
- “incorrectly classified”: The wrong label is assigned. For example, whisker C2 is correctly traced, but is labeled as C1. If the whisker is both poorly traced and incorrectly classified, it is labeled as incorrectly classified.
- “false positive”: A non-whisker object is reported as a whisker, or the same whisker object is labeled as two different whiskers.
- “false negative”: The whisker is present in the frame, but is not labeled. This is possible because our algorithm applies a confidence and smoothness threshold to the output of the neural network, and outputs that do not pass these thresholds are simply dropped.

We used an iterative procedure to train our algorithm, which was critical to its success.

1. First, we chose n = 7433 frames randomly from all sessions for which we had video, applied a previous generation whisker tracking algorithm (`whisk`), and manually labeled the identity of each whisker after verifying that it was correctly traced. This is “curated dataset 1”, representative of typical frames in the video.
2. We trained a neural network on that dataset, and used it to label all the frames in all the videos.
3. Of all these frames, we chose the frames on which the network was most likely to have made mistakes. We did this in several parallel ways: identifying frames where the reported confidence values were intermediate (i.e., unsure of presence or absence of the whisker), where the whiskers were near the extreme ends of their typical ranges, where any whisker was missing, and when any whisker was missing during a contact event. These are “challenge frames”, because they were chosen for their difficulty.
4. We manually evaluated and corrected every challenge frame, using the previous generation whisker tracking algorithm as a backup method when necessary.
5. We repeated steps 2-4 four times, to generate curated datasets 2-5.

As reported previously^6^, each type of error is nearly zero (<0.2%) on dataset 1, which is the only dataset representative of typical frames (Figure 2). Paradoxically, error rates increase with each subsequent set of challenge frames, because as the algorithm improves, the frames on which it still makes errors become more and more difficult. The most common type of error is the false negative, because we used a relatively strict confidence threshold. However, false negatives are also the least problematic, because we interpolated missing whiskers over frames.

In all cases we used doubly held-out cross validation: splitting the data into train, tune, and test sets. The network was always trained on the training set. With the tuning set, we used a grid search to identify the best choice of hyperparameters (pos_dist_thresh=17, locref_loss_weight=.01, confidence threshold=0.9, whisker smoothness threshold, number of training iterations=1e6, and in pilot experiments we selected global_scale as 1.0, jittered from 0.5 to 1.5). Then we used the test set to evaluate the performance using the hyperparameters chosen from the tuning set. This entire process was repeated three times so that every frame was in the train, tune, and test set exactly once.

### Alignment data

We assessed the quality of the synchronization between data streams by fitting a line to the timestamps of the synchronization pulses between any two time bases (e.g., behavior vs video, or video vs neural). The residuals around this linear fit reflected its accuracy. We observed that the behavioral timebase was a poor fit to the other two timebases and that its residuals varied slowly but predictably over the session, suggesting that the Arduino’s clock varied in rate with a timecourse of minutes. Therefore we used “trial by trial” alignment for the behavioral data such as lick times.

In contrast the neural and video timebase were synchronized very well by a linear fit. Residuals were never more than a single frame (5 ms) and thus the fit was essentially optimal given the resolution of the video data. Thus, we were able to apply this linear fit to the timestamps of the synchronization pulses in the neural data to predict the timestamps of the same pulses in the video data with sub-frame accuracy, as part of the “continuous alignment” procedure.

### Inclusion criteria

We excluded data from mice who were using a modality other than the whiskers to solve the task, which we assessed by trimming all of the whiskers after the last behavioral session. If the mouse performed significantly above chance on that day, indicating that it must be using another modality such as vision or olfaction to identify the objects, then we discarded all of the data from that mouse. We also excluded sessions with fewer than 90 trials to analyze.

As reported previously^6^, sessions with inaccurate labeling were discarded: we required that every whisker be labeled in ≥95% of the frames, that ≤2% of the contact events contained even a single frame with a missing label, and that the arcs traced out over the entire session by the whisker contained no discontinuities or jumps suggestive of tracking errors. In the remaining well-traced sessions we interpolated whiskers over any missing frames.

## Usage Notes

Users can load and analyze the data directly on hub.dandiarchive.org without downloading any files or installing any software locally. Alternatively, users can download any or all of the NWB files and analyze them locally using the PyNWB module. The MatNWB interface should provide equivalent access using Matlab, but we did not test it.

In this section, we provide some example code demonstrating how to load the data. It can be run directly on Dandi Hub without downloading any data. The same code is also available as a JupyterLab notebook at https://github.com/cxrodgers/NwbDandiData2022/blob/main/UsageNotes.ipynb.

First, we import the required modules.

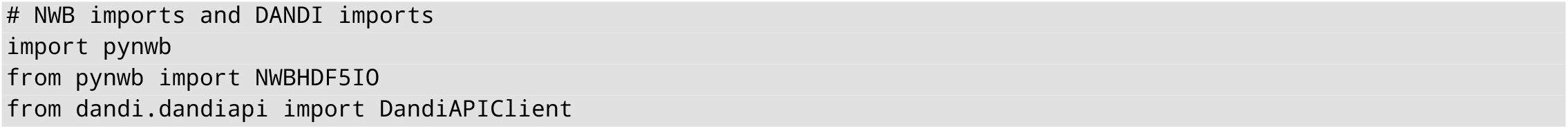

We specify the location of an example NWB file.

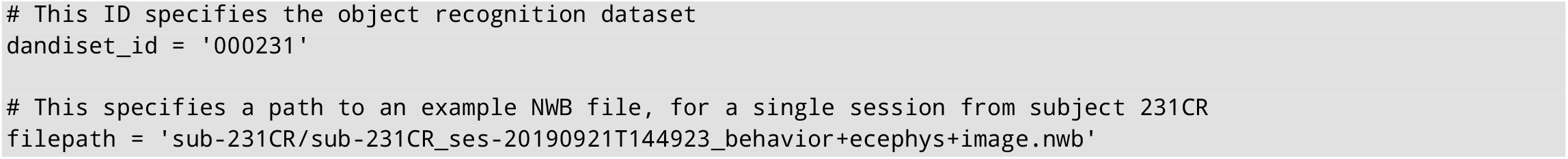

We load the data directly from DANDI.

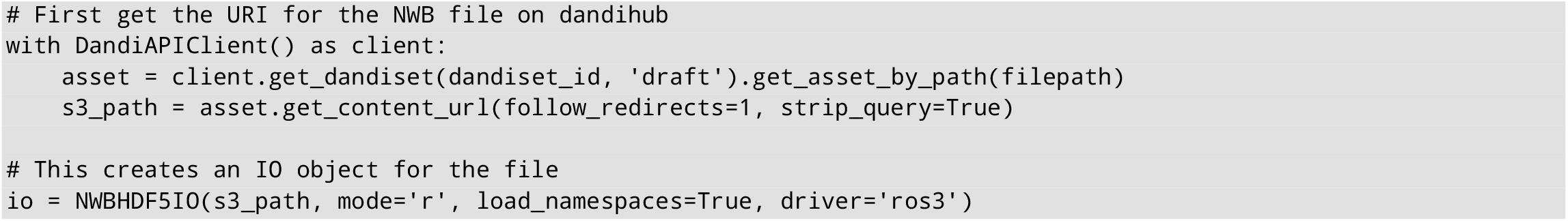

Alternatively, users could download the file(s) directly from DANDI, and specify the path to the downloaded NWB file, instead of the `s3_path` given above. In that case, there is no need to specify the “driver”. In any case, the next line reads the file into a variable called `nwbfile`, which we use for the rest of the demonstration.

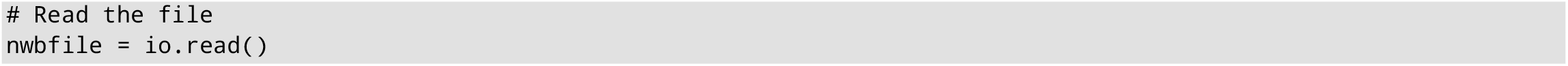

For the rest of the example code, the string “In [0]:” indicates a prompt, and the rest of the line following the prompt is the command the user should type. The lines following the prompt show the expected output.

It is often useful to first identify the time within the session at which each trial occurred, using the `trials` table.

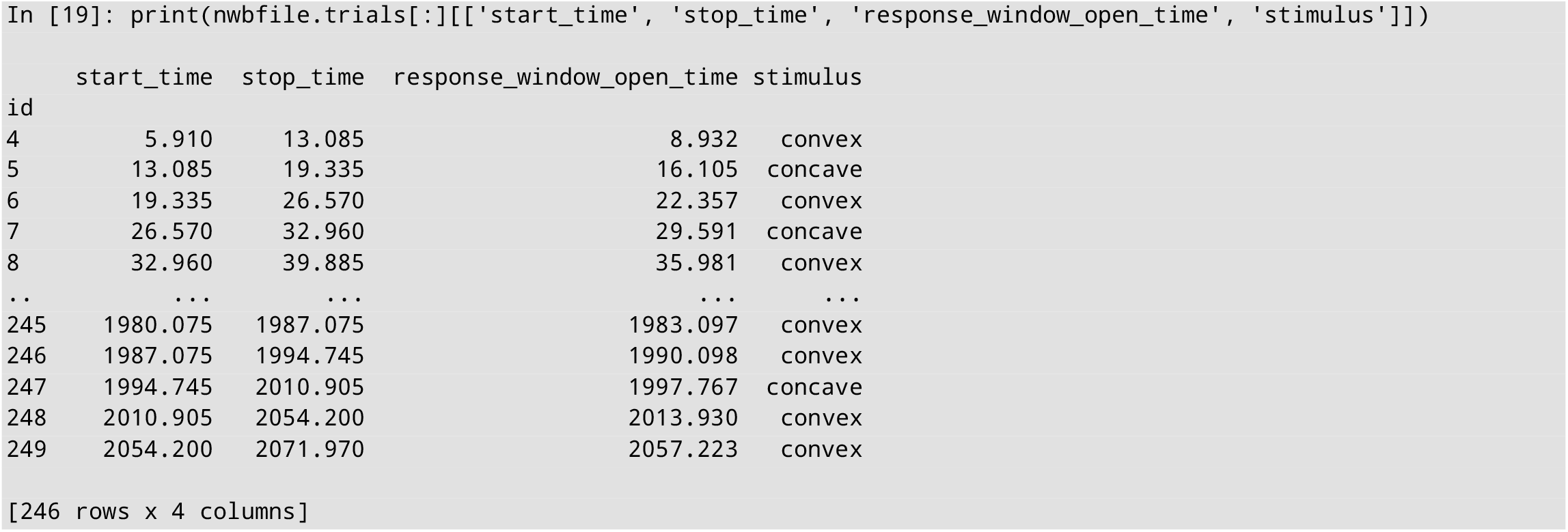

We suggest aligning data to the “response_window_open_time” on each trial, which is locked to the presentation of the shape stimulus (see “Methods” / “Session control”).

The position of the whiskers can be extracted over the entire session. Here we extract the angle of the C2 whisker over the entire session. The angle itself is stored in the field named `data`, and the timestamps of the angle are stored in the field name `timestamps`. Other field specify metadata.

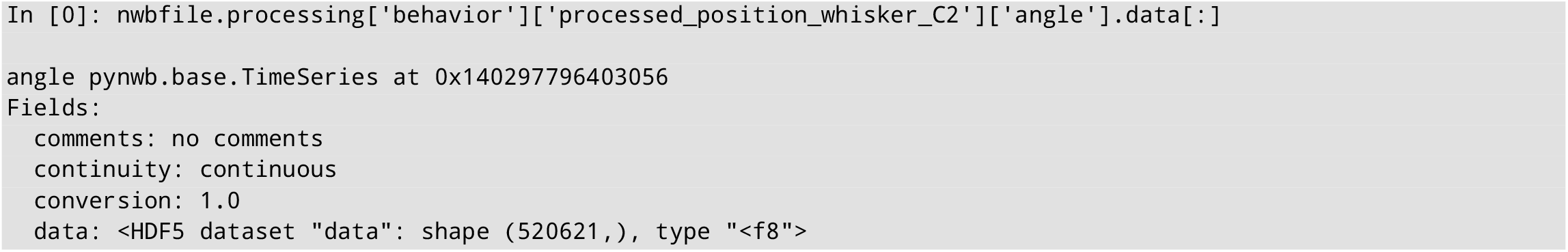
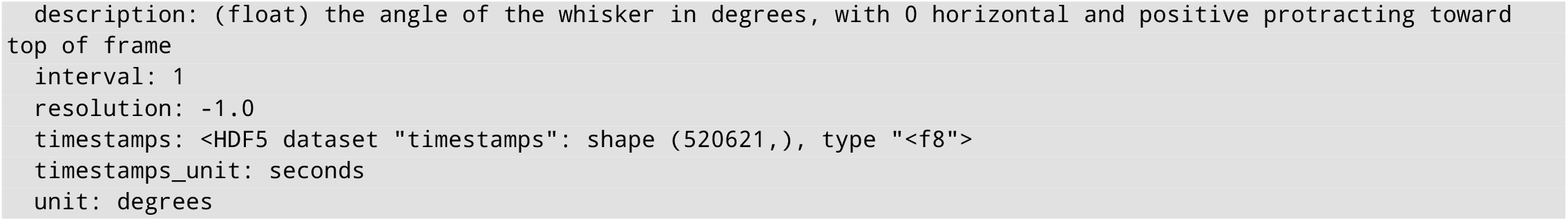

We can also extract the time of the contacts made by the same whisker.

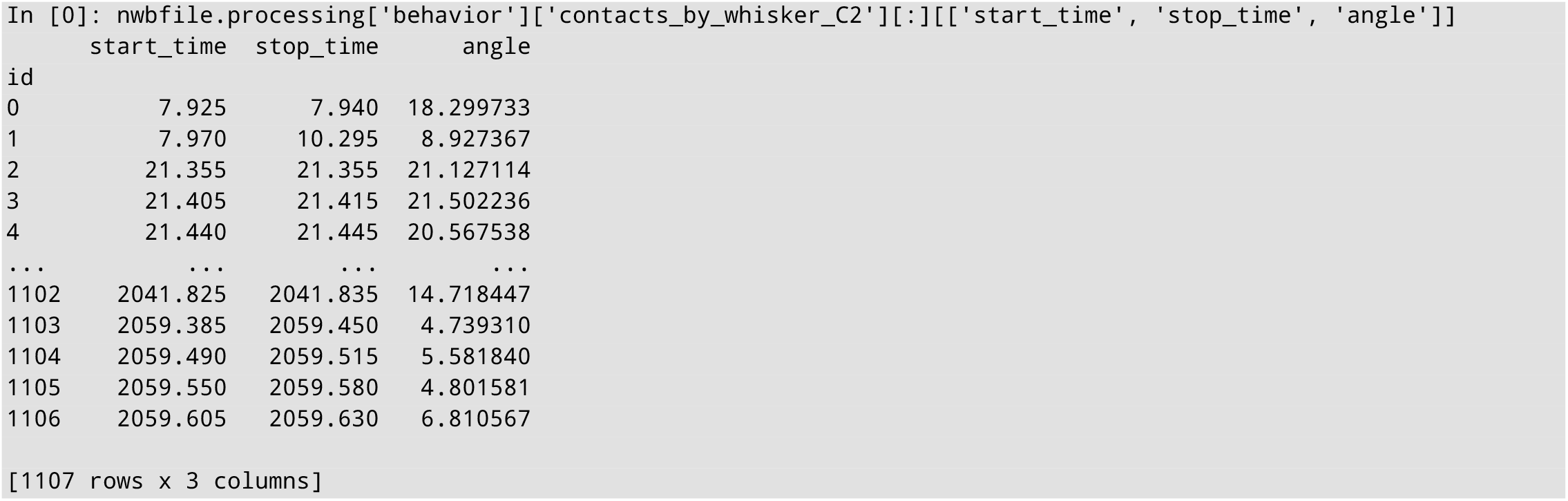

We can also extract full-bandwidth neural data from the entire session. Here, only the first 1e6 datapoints are loaded.

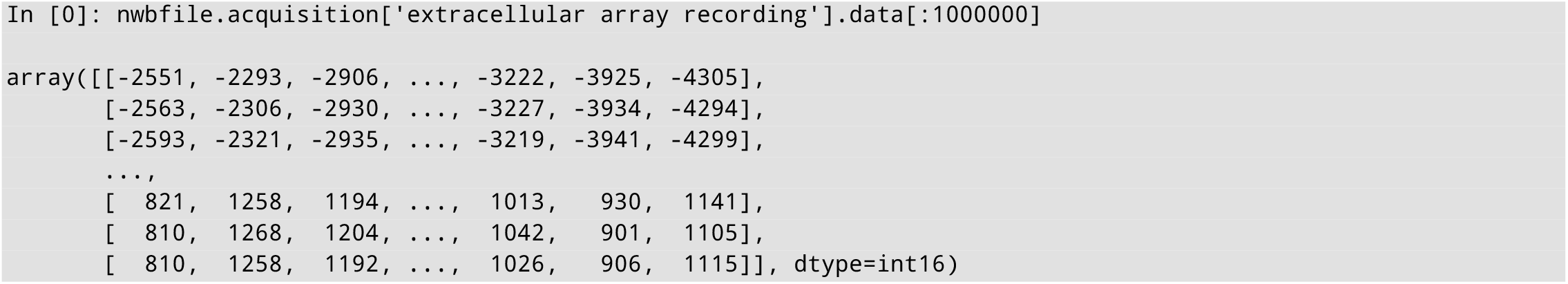

The `units` table specifies all the individual neurons that were recorded, and all of their identified spike times.

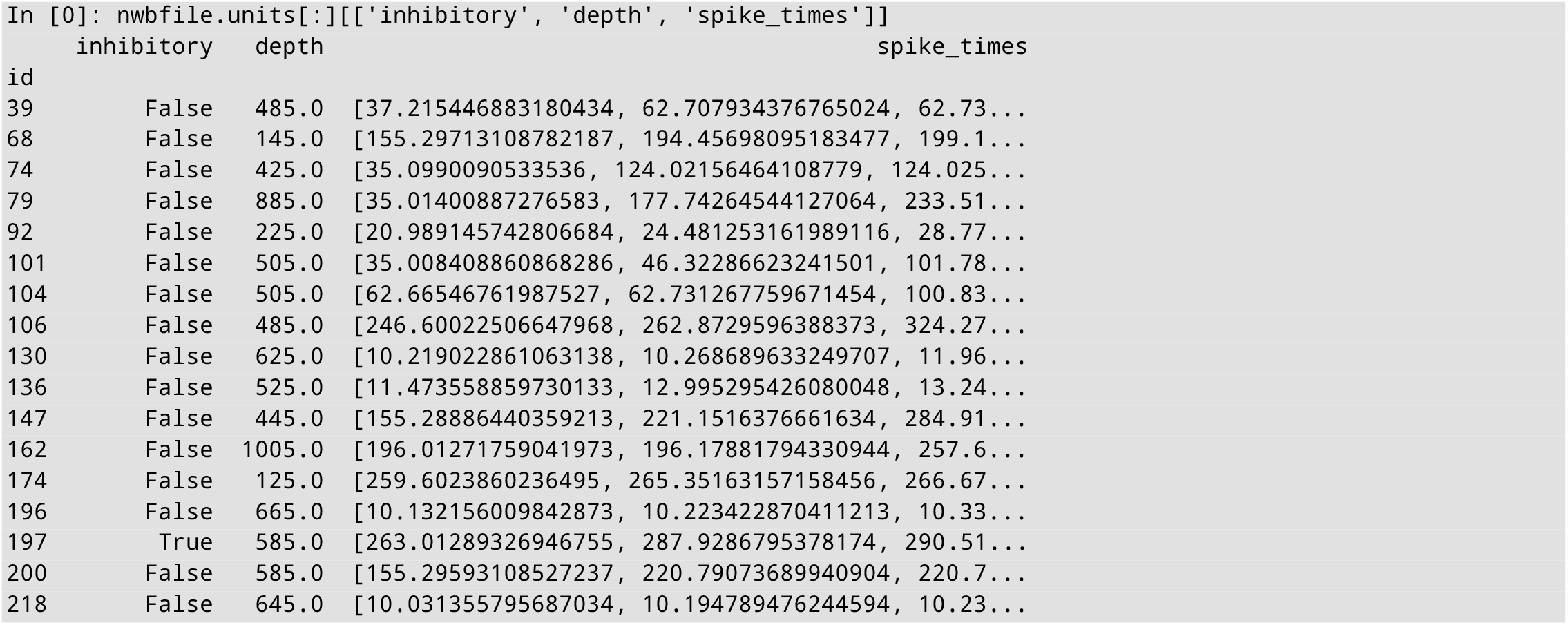

## Code Availability

We used the following packages to generate this dataset:

- Open Ephys acquisition GUI^16^, version 0.4
- Kilosort^18^, version 1
- Phy^19^, version 2.0a1
- pynwb^9^, version 2.0.0
- ndx_pose, https://github.com/rly/ndx-pose
- pose-tensorflow^12,13^, https://github.com/cxrodgers/PoseTF, lightly modified fork of https://github.com/eldar/pose-tensorflow
- whisk^11^, https://github.com/cxrodgers/whisk, lightly modified fork of https://github.com/nclack/whisk
- ffmpeg-python, https://github.com/kkroening/ffmpeg-python
- ffmpeg, version 4.0.2
- ipython^20^, version 7.22.0
- pandas^21^, version 1.2.4
- numpy^22^, version 1.19.2
- scipy^23^, version 1.6.2
- scikit-learn^24^, version 0.24.1
- scikit-image^25^, version 0.18.1
- statsmodels, version 0.12.2
- pyglmnet^26^, https://github.com/glm-tools/pyglmnet
- matplotlib^27^, version 3.3.4

## Acknowledgements

We would like to acknowledge Ramon Nogueira, B Christina Pil, Esther A Greeman, Jung M Park, Y Kate Hong, Stefano Fusi, and Randy M Bruno, who co-authored with CR the original paper based on this dataset. We would also like to acknowledge the members of the Neurodata Without Borders Slack and the DANDI Slack for their advice on using these data standards. The present work was supported in part by a Kavli Neurodata Without Borders Seed Grant (S-2021-GR-040) to CR. The original data collection was supported in part by a Kavli Institute Postdoctoral Fellowship, a NINDS/NIH NRSA fellowship (F32NS096819), and a NARSAD Young Investigator Award (28667) from the Brain & Behavior Research Foundation to CR.

## Author Contributions

CR collected the original dataset, wrote the methods for iteratively improving the pose tracking training procedure and for aligning the trials, converted the files to the NWB standard, and wrote the manuscript.

## References

1. Yang, S. C. H., Wolpert, D. M. & Lengyel, M. Theoretical perspectives on active sensing. Curr. Opin. Behav. Sci. 11, 100–108 (2016).

2. Gibson, J. J. The ecological approach to visual perception. (1979).

3. Grant, R. A., Breakell, V. & Prescott, T. J. Whisker touch sensing guides locomotion in small, quadrupedal mammals. Proc. R. Soc. B Biol. Sci. 285, (2018).

4. Lederman, S. J. & Klatzky, R. L. Hand movements: a window into haptic object recognition. Cogn. Psychol. 19, 342–368 (1987).

5. Yau, J. M., Kim, S. S., Thakur, P. H. & Bensmaia, S. J. Feeling form: The neural basis of haptic shape perception. J. Neurophysiol. 115, 631–642 (2016).

6. Rodgers, C. C., Nogueira, R., Pil, B. C., Greeman, E. A., Park, J. M., Hong, Y. K., Fusi, S. & Bruno, R. M. Sensorimotor strategies and neuronal representations for shape discrimination. Neuron 109, 2308-2325.e10 (2021).

7. Bale, M. R. & Maravall, M. Organization of sensory feature selectivity in the whisker system. Neuroscience 368, 70–80 (2018).

8. Nogueira, R., Rodgers, C. C., Bruno, R. M. & Fusi, S. The geometry of cortical representations of touch in rodents. bioRxiv 2021.02.11.430704 (2021). at <https://doi.org/10.1101/2021.02.11.430704>

9. Rübel, O. et al. NWB:N 2.0: An Accessible Data Standard for Neurophysiology. bioRxiv 523035 (2019). at <https://www.biorxiv.org/content/10.1101/523035v1>

10. Rodgers, C. C. Dataset of behavior and neural responses during shape discrimination and detection. Retrieved from https://dx.doi.org/10.5281/zenodo.4 (2021). xat <https://dx.doi.org/10.5281/zenodo.4743837>

11. Clack, N. G., O’Connor, D. H., Huber, D., Petreanu, L., Hires, A., Peron, S., Svoboda, K. & Myers, E. W. Automated tracking of whiskers in videos of head fixed rodents. PLoS Comput. Biol. 8, e1002591 (2012).

12. Insafutdinov, E., Pishchulin, L., Andres, B., Andriluka, M. & Schiele, B. DeeperCut: A Deeper, Stronger, and Faster Multi-Person Pose Estimation Model. Eur. Conf. Comput. Vis. 34–50 (2016). doi:10.1007/978-3-319-46466-4_3

13. Pishchulin, L., Insafutdinov, E., Tang, S., Andres, B., Andriluka, M., Gehler, P. & Schiele, B. DeepCut: Joint Subset Partition and Labeling for Multi Person Pose Estimation. (2015). doi:10.1109/CVPR.2016.533

14. Mathis, A., Mamidanna, P., Cury, K. M., Abe, T., Murthy, V. N., Mathis, M. W. & Bethge, M. DeepLabCut: markerless pose estimation of user-defined body parts with deep learning. Nat. Neurosci. 21, 1281–1289 (2018).

15. Stüttgen, M. C. & Schwarz, C. Barrel cortex: What is it good for? Neuroscience 368, 3–16 (2018).

16. Siegle, J. H., López, A. C., Patel, Y. A., Abramov, K., Ohayon, S. & Voigts, J. Open Ephys: An open-source, plugin-based platform for multichannel electrophysiology. J. Neural Eng. 14, (2017).

17. Hooks, B. M., Hires, S. A., Zhang, Y.-X., Huber, D., Petreanu, L., Svoboda, K. & Shepherd, G. M. G. Laminar analysis of excitatory local circuits in vibrissal motor and sensory cortical areas. PLoS Biol. 9, e1000572 (2011).

18. Pachitariu, M., Steinmetz, N., Kadir, S., Carandini, M. & Harris, K. D. Kilosort: realtime spike-sorting for extracellular electrophysiology with hundreds of channels. bioRxiv 061481 (2016). doi:10.1101/061481

19. Rossant, C., Kadir, S. N., Goodman, D. F. M., Schulman, J., Hunter, M. L. D., Saleem, A. B., Grosmark, A., Belluscio, M., Denfield, G. H., Ecker, A. S., Tolias, A. S., Solomon, S., Buzski, G., Carandini, M. & Harris, K. D. Spike sorting for large, dense electrode arrays. Nat. Neurosci. 19, 634–641 (2016).

20. Perez, F. & Granger, B. E. IPython: A System for Interactive Scientific Computing. Comput. Sci. Eng. 9, (2007).

21. McKinney, W. Data structures for statistical computing in Python. Proc. 9th Python Sci. Conf. (2010). doi:10.3828/ajfs.41.3.62

22. Van Der Walt, S., Colbert, S. C. & Varoquaux, G. The NumPy array: A structure for efficient numerical computation. Comput. Sci. Eng. 13, 22–30 (2011).

23. Virtanen, P. et al. SciPy 1.0: fundamental algorithms for scientific computing in Python. Nat. Methods 17, (2020).

24. Pedregosa, F., Varoquaux, G., Gramfort, A., Michel, V., Thirion, B., Grisel, O., Blondel, M., Prettenhofer, P., Weiss, R., Dubourg, V., Vanderplas, J., Passos, A., Cornapeau, D., Brucher, M., Perrot, M. & Duchesnay, É. Scikit-learn. J. Mach. Learn. Res. 12, 2825–2830 (2011).

25. Van Der Walt, S., Schönberger, J. L., Nunez-Iglesias, J., Boulogne, F., Warner, J. D., Yager, N., Gouillart, E. & Yu, T. Scikit-image: Image processing in python. PeerJ 2014, 1–18 (2014).

26. Jas, M. et al. Pyglmnet: Python implementation of elastic-net regularized generalized linear models. J. Open Source Softw. 5, 1959 (2020).

27. Hunter, J. D. Matplotlib: a 2d graphics environment. Comput. Sci. Eng. 9, 90–95 (2007).

